# Delayed formation of neural representations of space in aged mice

**DOI:** 10.1101/2023.03.03.531021

**Authors:** Kelsey D. McDermott, M. Agustina Frechou, Jake T. Jordan, Sunaina S. Martin, J. Tiago Gonçalves

## Abstract

Aging is associated with cognitive deficits, with spatial memory being very susceptible to decline. The hippocampal dentate gyrus (DG) is important for processing spatial information in the brain and is particularly vulnerable to aging, yet its sparse activity has led to difficulties in assessing changes in this area. Using *in vivo* two-photon calcium imaging, we compared DG neuronal activity and representations of space in young and aged mice walking on an unfamiliar treadmill. We found that calcium activity was significantly higher and less tuned to location in aged mice, resulting in decreased spatial information encoded in the DG. However, with repeated exposure to the same treadmill, both spatial tuning and information levels in aged mice became similar to young mice, while activity remained elevated. Our results show that spatial representations of novel environments are impaired in the aged hippocampus and gradually improve with increased familiarity. Moreover, while the aged DG is hyperexcitable, this does not disrupt neural representations of familiar environments.

## Introduction

As the world’s population ages, it becomes more important to understand the cognitive effects of aging and to develop therapeutic strategies for the aging brain. While healthy aging has milder cognitive effects than neurodegenerative states, several aspects of cognitive function exhibit decline (*1*). Memory deteriorates with age, with spatial memory being particularly susceptible to deficits in older humans and animals (*2*–*4*). Older adults show orientation deficits (*5, 6*), although they tend to perform better in familiar locations or tasks than in novel ones (*7*–*9*). This aging-related decay is thought to be caused by vulnerabilities in the hippocampal circuits that mediate spatial memory (*10, 11*). The dentate gyrus (DG) is one of the hippocampal subregions that are most vulnerable to aging (*12*–*14*), and the inputs from the entorhinal cortex to DG are a main area of dysfunction in both Alzheimer’s Disease (AD) patients and AD mouse models (*15*). During aging, there are reduced synaptic contacts from both the medial and lateral entorhinal cortex (*16*) onto DG neurons, as well as reduced synaptic plasticity of these inputs (*17*). However, the is very limited data about how DG activity changes with aging, primarily because neuronal activity in this region tends to be too sparse for *in vivo* electrophysiological recordings (*18*).

Here we investigated whether aging is associated with impaired DG activity and spatial representations, which could account for the spatial memory deficits seen in aged individuals. Since the DG is upstream of other hippocampal areas, changes in this region could help elucidate the causes of aging-related changes in neuronal activity (*19*–*22*) and neural representations of space (*19, 23, 24*) found in some areas of the hippocampus but not in others. Using *in vivo* two-photon microscopy we imaged calcium activity in a large population of DG neurons of young and aged mice as the animals walked head-fixed on a treadmill to which they had never been exposed. We found that aged mice have increased DG single-cell calcium activity and disrupted neural representations of space upon their first exposure to the treadmill setting. However, further imaging sessions on subsequent days showed that spatial representations become similar to those of young mice as the animals familiarize themselves with the treadmill, whereas single-cell activity remains elevated. Our results highlight the importance of novelty and familiarity to spatial encoding in aged animals.

## Results

To record neuronal activity in DG excitatory neurons, we injected young (3-4 month old) and aged (21-26 month old) C57Bl6 mice with an AAV encoding the calcium indicator jRGECO1a (*25*) under the CaMKII promoter (Fig. 1A). We then implanted the mice with a hippocampal imaging window above the DG (*26, 27*) to enable *in vivo* two-photon calcium imaging in this region (Fig. 1A,B). After waiting 3-4 weeks to allow for viral expression and recovery from surgery, we imaged mice on four consecutive days. During each imaging session the mice ran head-fixed along a treadmill containing tactile cues with four distinct segments (Fig. 1A,C) that was manually moved (Fig. S1). We recorded 10 minute videos of calcium activity and identified the regions corresponding to active cells using Suite2p software (Fig. 1D). Previous studies found that hippocampal activity in area CA3 becomes elevated during aging (*19, 28*). We determined single-cell calcium activity by measuring the area under the curve of transients and normalizing to the distance run by each mouse (*29*) (Fig. 1E-G). We found that the aged group had higher activity levels than the young group (Fig. 1E, p=0.00463, nested bootstrap), which aligns with previous reports of hyperactivity. We also estimated the fraction of active neurons during each imaging session and found no changes between groups (Fig. S2). We next investigated whether there were differences between the spatial representations of young and aged DG neurons. Both young and aged DG neurons have a variety of spatial responses, including place cells that only respond to a specific location and neurons whose activity shows low spatial preference (Fig. 2A,B). We determined the spatial selectivity of neuronal activity using a previously described spatial tuning index (see methods). DG neurons in aged mice had significantly lower tuning indices than those in young mice (Fig. 2C, p<0.0001, nested bootstrap). This difference indicates that the activity of DG neurons in aged mice was not as place-specific as that of young DG neurons, suggesting an aging-related degradation of the spatial code. We also computed single-cell Fisher Information (FI), as a measure of how much spatial information was encoded by individual DG neurons. By plotting whole-recording calcium activity raster plots for neurons in the 10^th^ percentile of FI we verified that neurons with high spatial information behave like ’place cells’ (*30*) in both young and aged animals, as their activity is concentrated at a single location on the treadmill (Fig. 2D,E). This place-specific response was present across laps as neurons in both cohorts of mice largely kept the same place response in both even and odd laps. Aged mice encoded less spatial information than young mice as denoted by a significant decrease in FI (Fig. 2F, p=0.00005, nested bootstrap).

**Figure 1.**
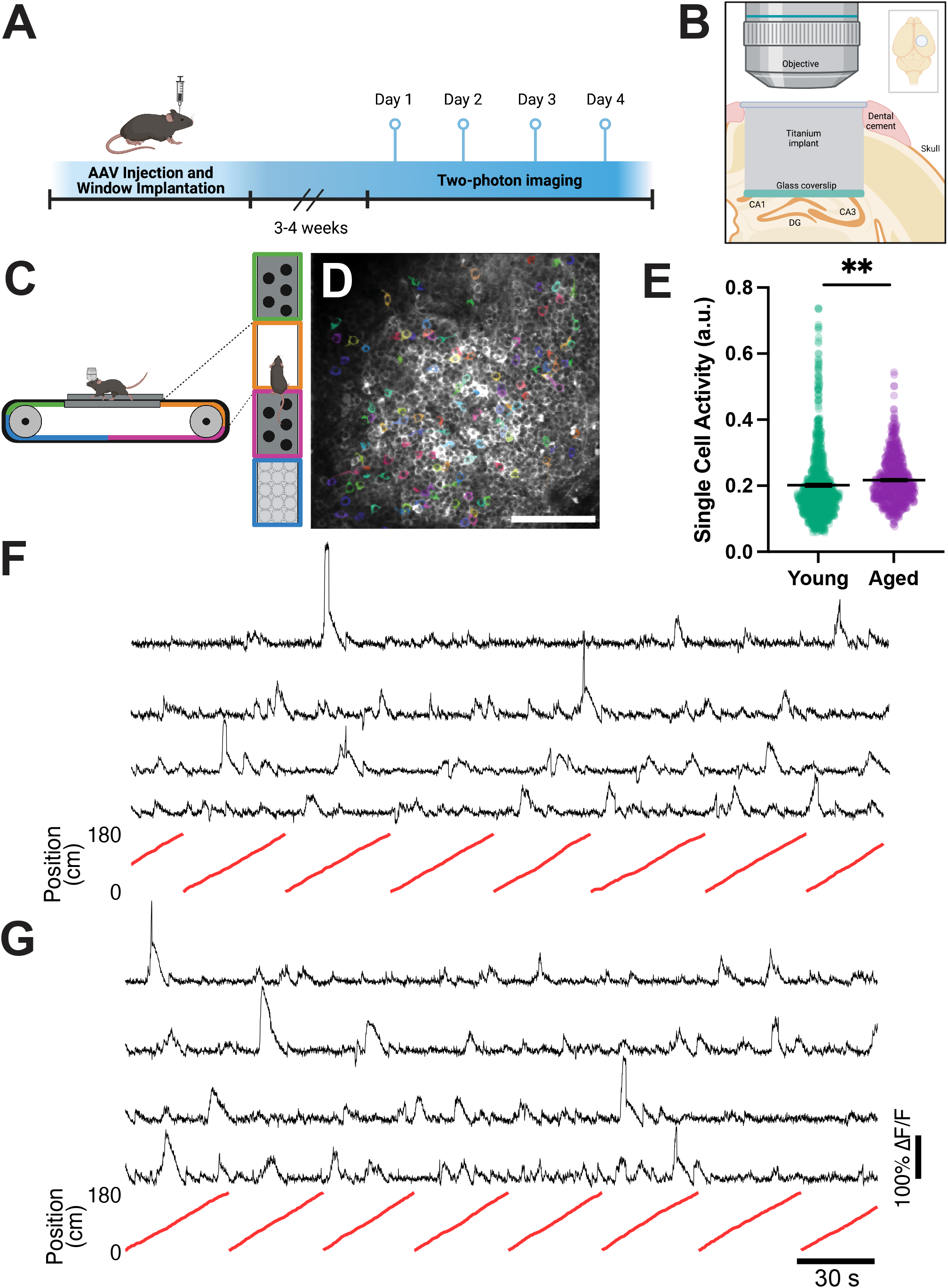
DG is hyperactive in aged animals. A) Experimental timeline, including AAV injection, window implantation, and two-photon imaging. B) Diagram of chronic window implant over the right hemisphere of the DG. C) Diagram of imaging setup, with side view (left) and top view (right) of mouse head-fixed to treadmill with multiple tactile zones. D) Example field of view with regions of interest of active cells shaded in color. Scale bar = 100 μm. E) Mean single cell calcium activity.F)Example calcium traces (black) and corresponding treadmill positions (red) from young mice.G)Example calcium traces (black) and corresponding treadmill positions (red) from aged mice. a.u.= arbitrary units. Young N=8 mice, n=910 cells; Aged N=8 mice, n=699 cells. **p<0.01

**Figure 2.**
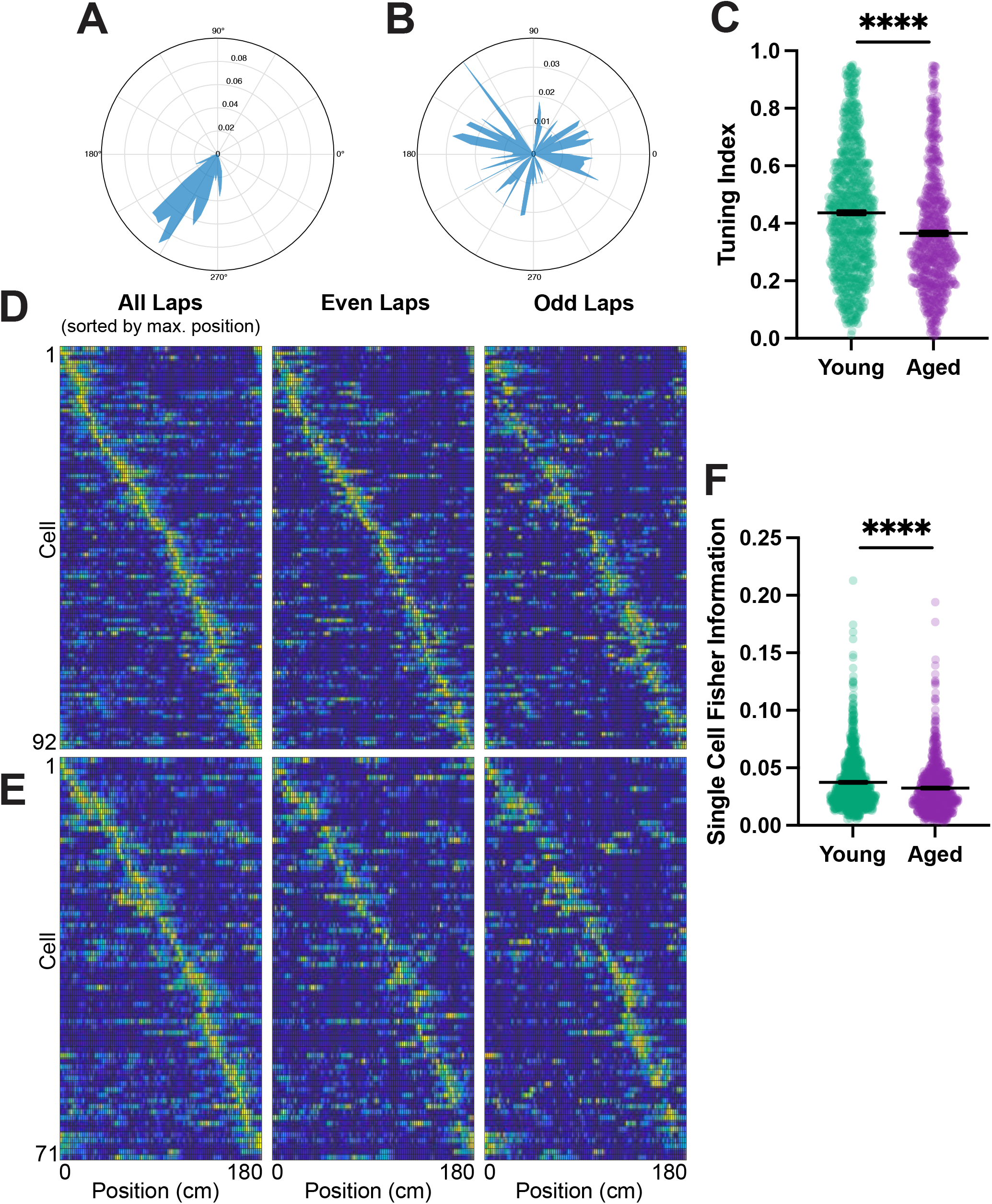
DG representations of space are impaired in aged mice. A) Tuning vector polar plots representing a highly spatially tuned cell and B) a cell with low spatial tuning. C) Spatial tuning index. D) Raster plots of tuning vectors of young mouse neurons in 10^th^ percentile Fisher Information, sorted by the position of activity maximum for whole recording (All Laps). Raster plots of partial recording (Even Laps and Odd Laps) keep same sorting used for whole recording. E) Same raster plots for aged mice. C) Single cell Fisher information. Statistics done with nested bootstrap analysis. Bars represent mean +/- SEM. a.u.= arbitrary units. Young N=8 mice, n=910 cells; Aged N=8 mice, n=699 cells. *p<0.05, **p<0.01, ***p<0.001, ****p<0.0001. See also supplemental figures 1 and 2.

To confirm that the changes in spatial representations in aged mice corresponded to a change in spatial memory, mice underwent behavioral testing using an object placement paradigm. This test takes advantage of a mouse’s natural preference for novelty to assess how well animals can remember the location of objects in space, a behavior that is thought to be hippocampus-dependent (*31*). We allowed mice to explore an arena with visual cues and two novel and identical objects during a training trial. We then displaced one object to another location in the arena and again allowed the mice to explore during a test trial (Fig. S3A). Young mice had a preference for the novel object that was significantly above chance (p= 0.0017, One sample Wilcoxon test) whereas aged mice did not score above chance (Fig S3B-C, p=0.6355, One sample Wilcoxon test). Additionally, significantly more aged mice than young mice failed the test (Fig. S3D, p=0.0393, Chi squared test). This change in preference can also be seen in the exploration times per objects, in which young mice spend more time engaged with the displaced object, while aged mice spend about an even amount of time between objects (Fig. S3E, p=0.0205, Two-way ANOVA with Sidak multiple comparisons test). These data confirm earlier findings of spatial memory deficits in aged mice, which we now show is accompanied by impaired neural representations of space in the DG.

Older individuals are better able to distinguish places in an environment that they are familiar with than in a novel environment (*7, 8*), so we asked how the representations of space in the DG changed across four consecutive days of imaging, as the treadmill belt became more familiar to the mice. Single-cell calcium activity remained elevated in aged mice through all recording days, even though this difference was not significant in days 2 and 3 (Fig. 3A, nested bootstrap with Bonferroni correction). While in young mice there was a slight but significant decrease in activity (p=0.00005) across days, in aged mice activity increased over time (p<0.0001) (Fig. 3B-C, nested bootstrap). In contrast to activity, the differences in tuning index and FI were erased over the four days of imaging, as young and aged groups converged. The spatial tuning index of young mice remained approximately constant over the course of all imaging sessions (p=0.19772), whereas the tuning of aged mice underwent an increase so that the differences between both groups were no longer significant in days 3 and 4 (p=0.00005). (Fig. 3D-F, nested bootstrap, Bonferroni corrected in 3D). FI rose in both young and aged groups over the four imaging days, but the increase was more pronounced in aged animals so that the differences between young and aged animals were no longer significant in days 3 and 4 (Fig. 3G-I, nested bootstrap, Bonferroni corrected in 3G). These data suggest that representations of space in aged animals can be rescued by repeated exposure to the same environment.

**Figure 3.**
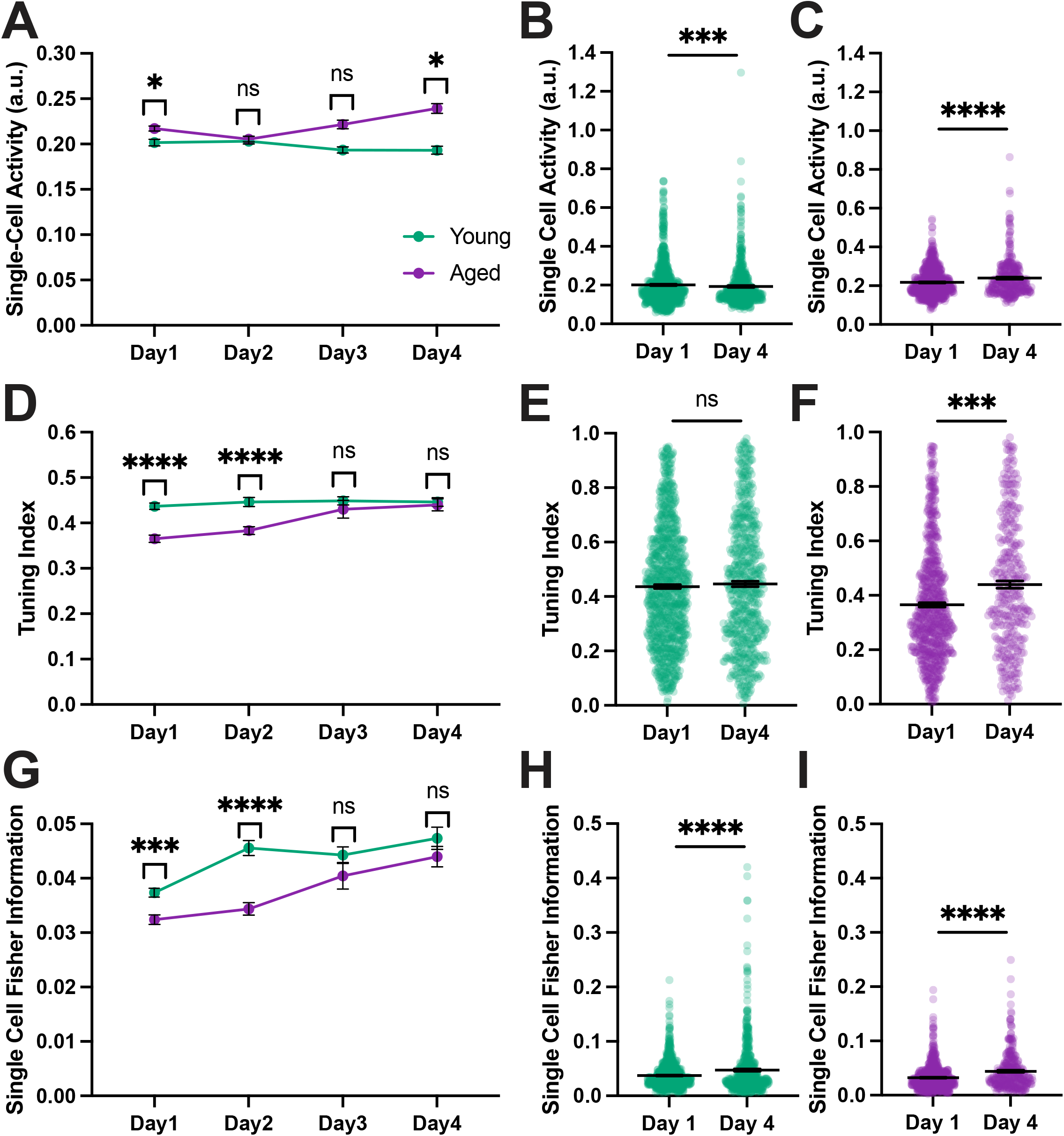
Deficits in aged spatial representations are rescued with increased familiarity. A) Mean single cell calcium activity across days. B) Single cell activity in young mice on day 1 versus day 4. C) Single cell activity in aged mice on day 1 versus day 4. D) Mean tuning index across days. E) Tuning index in young mice on day 1 versus day 4. F) Tuning index in aged mice on day 1 versus day 4. G) Mean single cell Fisher Information across days. H) Fisher Information in young mice on day 1 versus day 4. I) Fisher Information in aged mice on day 1 versus day 4. Statistics done with a nested bootstrap analysis. Bars represent mean +/- SEM. a.u.= arbitrary units. Young: day 1 N=8, n=910; day 2 N=7, n=694; day 3 N=7, n=623; day 4 N=7, n=610; Aged: day 1 N=8, n=699; day 2 N=8, n=619; day 3 N=5, n=147; day 4 N=6, n=331. ns=not significant, *p<0.05, **p<0.01, ***p<0.001, ****p<0.0001.

Previous studies found the stability of neural representations of space was altered in aged animals (*32, 33*). We therefore tracked the activity of individual neurons across days to investigate the stability of the active cell ensemble (Fig. 4A-C). We first asked whether cells that were active in the initial imaging session were reactivated on subsequent days. Surprisingly, we found no difference between groups, suggesting that both DG neurons in young and aged mice have similar reactivation rates (Fig. 4D, mixed effects model with Sidak multiple comparisons test). We went on to compare the activity in the cells that were active both on day 1, when the treadmill was novel, and day 4, when it had become familiar. The single-cell activity of matched cells increased across days in the young mice (p= 0.00234, nested bootstrap) but not in the old mice (p=0.15981, nested bootstrap) (Fig. 4E-F). This may reflect a ceiling for activity since the aged mice already had higher activity on day 1. While the tuning index did not differ in either group across days (Fig. 4G-H, p=0.07626, p=0.21341), both groups saw higher spatial information, as FI was significantly increased in reactivated cells on day 4 (Fig. 4I-J, p=0.00015, p=0.00024), which may simply reflect the fact that FI increases sharply for the entire population across the four days. We then asked whether the reactivated cells had a distinct profile compared to those neurons that were not active across days. In day 1, cells that were to be reactivated did not differ from other cells in single cell activity levels. Whether cells were reactivated or not, their activity levels were higher in the aged mice on day 1, which recapitulates the pattern seen in the general population (Fig. 4K). These effects were gone by day 4, again reflecting the trend in the general population (Fig. 4L). In the aged group, cells that were reactivated showed higher tuning levels than other cells on day 1 (Fig. 4M), but this effect was also absent on day 4 (Fig. 4N). Reactivated cells had significantly higher FI than other neurons in both the young and aged groups on both day 1 and day 4 (Fig. 4O-P). This suggests that higher spatial information is a predictor of whether neurons will be reactivated over several days. Overall, we did not see evidence any changes in the stability of coding ensembles in aged animals, when compared with the young cohort. However, since reactivated neurons initially have higher tuning and spatial information than non-reactivated cells in aged animals, and increase their FI over time, it’s possible that they contribute to the improvement of spatial representations in aged mice improve over the four days of imaging.

**Figure 4.**
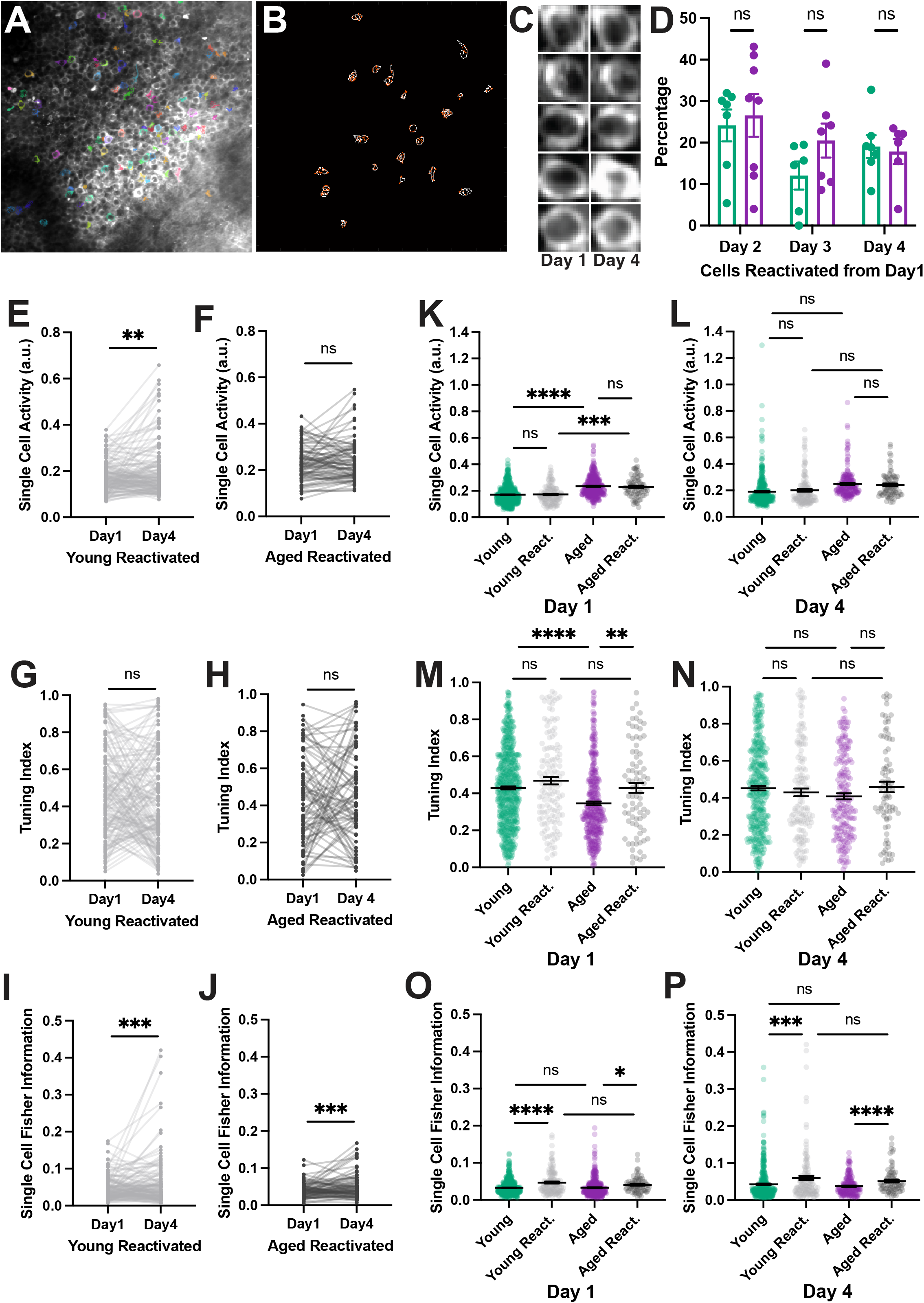
Reactivated cells do not have a distinct profile in aged animals. A) Example field of view used to match cells across days. Neurons that were active during recording session are shaded in color. B) Matched regions of interest from the field of view in panel A, red and white contours represent cells active on day 1 and 4. C) Examples of matched cells on day 1 and 4. D) Reactivation rate of neurons that were active on day 1. Percentages are based on imaging day versus day 1, regardless of whether the cell was active on other days. E-F) Matched single cell calcium activity in reactivated cells in young (E) and aged (F) mice. G-H) Matched tuning index in reactivated cells in young (G) and aged (H) mice. I-J) Matched single cell Fisher Information in reactivated cells in young (I) and aged (J) mice. K-L) Single cell activity of reactivated and non-reactivated cells on day 1 (K) and day 4 (L). M-N) Tuning of reactivated and non-reactivated cells on day 1 (M) and day 4 (N). O-P) Single cell Fisher Information of reactivated and non-reactivated cells on day 1 (O) and day 4 (P). Statistics done with nested bootstrap analysis. Bars represent mean +/- SEM. a.u.= arbitrary units. Young: N=6; day 1 n=617; day 1 react. n=140; day 4 n=396; day 4 react. n=140; Aged: N=5; day 1 n=375; day 1 react. n=80; day 4 n=196; day 4 react. n=80. ns=not significant, *p<0.05, **p<0.01, ***p<0.001, ****p<0.0001.

In summary, we have found that, in aged animals, DG neurons are hyperexcitable and impaired in their encoding of spatial features in novel environments. Our results highlight how the defects in spatial representations in the DG are specific to the first introduction to a novel context and can be rescued by increasing familiarity with a given environment.

## Discussion

Aging is associated with cognitive decline in spatial memory in healthy aged humans and rodents (*2*–*4*). In this study we asked whether aged mice have impaired representations of space in the DG circuits that mediate spatial memory, which could account for the memory deficits seen in aged individuals.

Several studies have found that neuronal activity is elevated in the hippocampus of aged animals specifically in hippocampal area CA3 (*19*–*22*). Human studies using fMRI in older and younger adults have generally confirmed the hyperexcitability findings in the DG-CA3 axis (*5, 14*), although fMRI does not have the resolution to distinguish between areas DG and CA3. This aging-related hyperexcitability phenotype is thought to contribute to the memory deficits seen in aged individuals (*34, 35*). In order to understand the causes of the aging-related hyperexcitability it would be important to record neuronal activity from the DG, as this region provides the main inputs to CA3.

We used *in vivo* two-photon calcium imaging to record activity from young and aged DG neurons. This allowed us to image a large population of cells and therefore have a better chance capturing active cells in a sparsely active population. Additionally, using an imaging-based approach allowed us to study the same group of cells across time. We have found that single-cell calcium activity is indeed increased in aged DG neurons, when compared to young controls (Fig. 1E).

We also found that during the initial exposure to the treadmill, there is a significant reduction in spatial information and tuning in aged DG neurons, as compared to their young counterparts (Fig. 2B-C). By using untrained mice for our experiments we were able to capture the very first neuronal representations of a novel environment. We found that during the course of four consecutive imaging days, spatial information and tuning in the aged mice converged with those of young mice, which leads us to conclude that aged mice require additional exposure to a novel environment in order to form adequate neural representations of space. It can therefore be said that the spatial encoding defects seen in aged mice are specific to novel environments. This is in agreement with previous reports that found that spatial representations in CA1 were disrupted when novel environmental cues were introduced during a navigation task (*33, 36*). Interestingly, both young and aged groups saw an increase in spatial information through the four days of imaging, whereas spatial tuning increased in aged mice but was not significantly different in young mice. We did not find a convergence in single-cell calcium activity levels between aged and young groups, meaning that the hyperexcitability is no impediment to aged mice eventually forming accurate spatial representations. There are several interesting parallels between our findings and human data showing that aged individuals are better able to navigate familiar environments than novel ones (*8*) leading them to avoid taking novel routes in favor of navigating familiar routes.

To verify that aged mice have impairments in spatial memory, we used an object placement test that is not physically tasking and does not involve an aversive component. Our behavior data confirmed that aged mice are more likely to fail at an object placement task than their younger counterparts. This goes along with studies analyzing similar spatial memory tasks that have found deficits in aged animals (*37, 38*). Our results also highlight the well-described heterogeneity present during normal aging, as about half of aged mice were still able to pass the test (*39, 40*).

In order to better compare our data to previous studies of DG activity using IEG labelling, we determined the percentage of active cells in our imaging fields but found no difference in the number of active cells between young and aged animals. This may be due to differences in the activation of ensembles based on the experimental designs. However, there are also some caveats to our experimental data that must be kept in mind: we calculate the total number of cells from mean projections of whole calcium imaging movies, which can introduce several biases, for example active cells will tend to be brighter and therefore more visible. We found a correlation between the percentage of active cells and the total number of cells in the imaging field (Fig. S2B), as fields of view with fewer detected cells had much higher variance of the fraction of active cells. Fields with more total cells generally had more cells that were dimmer and less active but were still counted in the total. In contrast, in fields with few cells due to areas of lower viral expression or obscured by blood vessels, there are likely more cells present than can be quantified in the mean projection image. These differences in imaging fields across animals makes it difficult to calculate exact percentages, limiting the usefulness of this approach.

Previous studies have found that increased hippocampal excitability contributes to aging-related cognitive impairments. Our experimental design did not allow us to determine whether hyperexcitability contributes to the deficits in spatial information seen in the novel environment during the first day of imaging, but our results suggest that instead dysfunction in the plasticity mechanisms that underlie spatial selectivity may be to blame. Spatial selectivity can develop very fast in CA1 hippocampal neurons through a mechanism termed behavioral timescale synaptic plasticity (*41*) that is mostly active in novel contexts, leading to the development and stabilization of spatial tuning within the first few minutes of exposure to a new environment (*42*). While less is known about the dynamics of spatial selectivity in the DG, spatially tuned cells also seem to emerge and stabilize within the first few minutes of exposure to a novel environment, with further refinement as the animals are re-exposed to the same environment over the following days (*43*). This is in line with our finding that spatial information increases over several days both in young and aged animals. Previous literature has shown that aging is associated with plasticity deficits in the hippocampus (*17, 44, 45*), namely in long term potentiation and depression. Given that these plasticity mechanisms are likely most active as mice map out new environments (*42*) we speculate that this may explain the protracted development of spatial selectivity that we see in aged mice. Expression of some IEGs, such as *Arc* and *Egr1*, which are thought to be regulators of neuronal plasticity, is reduced in the DG of aged animals. Deficits in the expression of these genes following neuronal activity could be at the origin of plasticity defects in aged animals. The increased excitability of aged DG neurons could therefore also be seen as a compensation mechanism for both the reduced synaptic input from the entorhinal cortex, and the reduced expression of IEGs. Another factor potentially contributing to the delayed formation of spatial representations in the aged DG is a reduction of adult neurogenesis. Adult-born neurons are known to enhance DG neuronal plasticity (*46, 47*) but their numbers fall to almost zero in aged animals (*48*).

Aging-associated memory deficits have also been associated with changes in the stability of spatial representations as some previous studies have found that the aged hippocampal spatial code undergoes larger changes over different sessions than in young mice (*32*), whereas others have found that spatial representations are more rigid (*24, 33*). We did not see changes in the reactivation of cells from one session to another between groups (Fig. 4D). This is in contrast to data showing that more distinct dentate populations express the IEG *Arc* when aged mice are re-exposed to the same environment (*49*), which could be due to age-related changes in DNA methylation rather than changes in neuronal excitability (*50*). In order to further investigate the properties of cells in the aged DG over time, we matched cells that were active in our imaging fields across days. We did not find major differences in the reactivated cells in any measures between young and aged groups, suggesting that ensembles of neurons follow the same basic mechanisms in the aged DG. When comparing reactivated neurons to the remainder of the population, we found that reactivated cells had significantly higher spatial information (Fig. 4 O,P), indicating that this may be a marker of whether a neuron will contribute to spatial representations over time.

To summarize, our data shows that new DG spatial representations are initially impaired and take longer to form in aged mice. These data expand our knowledge of the network activity and spatial representations in the aged hippocampus and suggest that aging-associated hippocampal hyperactivity is not an impediment to the formation of rich spatial representations. The protracted refinement of spatial representations suggests that the plasticity mechanisms responsible for the development of spatial selectivity are impaired in aged animals and are likely to be a relevant therapeutic target for ameliorating the memory deficits associated with aging.

## Acknowledgements

We thank Dr. Carolyn Pytte for the generous gift of aged mice from her laboratory at Queens College. We thank Dr. Maria Gulinello, the director of the behavior core facilities at Einstein, for her training and technical advice on behavior experiments. We thank Dr. Alberto Cruz-Martín for helpful feedback and discussions.

## Author contributions

Conceptualization: KDM, JTG; Methodology: KDM, MAF, JTJ, JTG; Investigation: KDM, SSM, JTG; Visualization: KDM, JTG; Supervision: JTG; Writing—original draft: KDM, JTG; Writing— review & editing: KDM, JTG

## Data availability

The analysis code used in this study is openly available at https://github.com/GoncalvesLab/McDermott-et-al-aging. All imaging data, consisting of calcium traces for every active cell and the position of the mouse on the treadmill, will be made openly available on a repository site before final publication. Due to their large size, raw calcium imaging movies will be made available upon request.

## Methods

### Animals

Aged mice were 21-26 months old C57Bl6J females originally from Jackson Labs and kindly gifted by Dr. Carolyn Pytte of Queens College. Young mice were 3-4 months old C57Bl6J females purchased from Jackson Labs. All mice were housed in standard conditions with a 14/10 hour light/dark cycle. Mice were provided food and water ad libitum. All procedures were done during the light part of the cycle and in accordance with the Einstein Institutional Animal Care and Use Committee (Protocol #00001197).

### Viral injections

Mice were anesthetized (induction: 5%, maintenance 0.5% isoflurane in O<sub>2</sub> vol/vol). Following anesthetization, mice were attached to a stereotactic apparatus and the right hemisphere of the dentate gyrus (DG) was injected with 1μl of a DJ-serotype AAV vector encoding the red-shifted calcium indicator jRGECO1a (*25*) under the CamKII promoter at 10^12^ GC/ml titer. Viral injections were done with a pulled glass pipette using a Nanoject III microinjector (Drummond) at previously described coordinates (*51*).

### Window implantation

Following viral injection, dexamethasone (1 mg/kg) was administered via subcutaneous injection to minimize brain swelling. Optibond (Kerr Dental) adhesive was applied to the skull and cured to help secure the implant once attached. A 3mm diameter craniotomy was made over the right dorsal DG and the overlaying tissue was removed by aspiration down to the hippocampal fissure where a custom-built cylindrical titanium implant with a glass coverslip on the bottom was inserted over the dorsal surface of the DG (*27*). The implant and a titanium bar for head-fixation were attached to the skull with dental cement. All mice were given carprofen (5 mg/kg, subcutaneous) as a post-surgery analgesic.

### Two-photon Imaging

*In vivo* calcium imaging movies were acquired with a custom two-photon laser scanning microscope (based on Thorlabs Bergamo) using a femtosecond-pulsed laser (Coherent Fidelity 2, 1070 nm) and a 16x water immersion objective (0.8 NA, Nikon). Imaging sessions were started 3-4 weeks after surgery to allow for recovery and for optimal viral expression. During imaging sessions, mice were head-fixed to the microscope with a titanium headbar and the microscope stage was adjusted so that the hippocampal window was perpendicular to the axis of the objective for optimal imaging conditions. Mice were awake and walking on a manually rotated treadmill belt throughout imaging. The treadmill contained four zones with different textures as previously described (*52*). Ten-minute videos were acquired at 15.253 fps with a 343.6 × 343.6 μm field of view. Treadmill data was acquired using National Instruments analog-to-digital converter and synchronized with the imaging data using Thorsync software. To track the same field of view, the initial location was noted on the first imaging day using a coordinate system and by taking an image of the field. During following sessions, the coordinates and initial field image were used to match as close as possible to the same field.

### Data Analysis

The open-source Suite2p software (version 10.0) was used to register and motion correct videos, for cell detection and spike deconvolution (*53*–*55*). Regions of interest corresponding to active neurons were identified by Suite2p and further manually curated in order to identify all active cells in the field of view. The ROIMatchPub (*56*) was used to identify and match cells in the same field of view across imaging sessions. Matched ROIs were manually confirmed after automated detection.

Single-cell calcium activity was determined as previously described (*29*). Briefly, ΔF/F data was filtered with a zero phase shift, third order Butterworth lowpass filter. This filtered ΔF/F was thresholded to 2 standard deviations(σ) above a rolling-mean baseline. Single cell activity was determined as the cumulative sum of the thresholded trace for each cell. This was then normalized to the total distance traveled by each mouse.

Tuning indices were using deconvolved calcium traces, which were thresholded to 2σ above the baseline. Any point not significantly above that noise threshold was set to zero. Videos were cut to only include periods of movement >1s in duration. The treadmill band was segmented into 100 position bins, and transients were mapped to these bins according to the location of the mice on the treadmill in order to generate a tuning vector for each cell (*26*).The mean of the thresholded activity at each location was calculated for every neuron and normalized to the time the mouse spent at that position. The tuning index was defined as the modulus of this normalized tuning vector.

Single-cell Fisher Information (FI) was calculated as previously described (*29, 57*). A bias-corrected signal to noise ratio was computed using the unfiltered ΔF/F fluorescence data for each individual cell, where the signal is the square of the difference of the mean activity at two locations on the treadmill, and the noise is the average variance of the activity at each location. Position data was segmented into 20 bins.

### Behavior

Object Placement –A square-shaped arena 42×42×30 (LxWxH) was set up with visual cues along the walls and with two novel and identical objects on the floor. Mice were individually placed in the arena for a 4 minute training trial. Following a 45 minute retention period, one object was displaced to another location in the arena and the mice were placed again for a 4 minute test trial. Exploration of the objects was quantified by an experimenter and confirmed with computer tracking.

### Statistics

Nested bootstrap analysis – When pooling cell data from different mice into a single experimental group, significance testing was done using a muti-level bootstrapped approach as previously described (*29*). To assess whether differences between two experimental groups are significant, a null surrogate distribution was constructed for each mouse in each group by resampling with substitution from the pool of all imaged cells. The difference between the null distributions generated for each group was calculated and the previous steps repeated to generate 100,000 bootstrap estimates of the difference between two groups’ null distributions. The empirically observed value of the difference between conditions was then compared to the null distribution values with a statistical significance level (α) set at 0.05. If the empirical group differences fall outside of the 95^th^ percentile of the 100,000 bootstrap estimates of the difference between the null distributions, then it is considered to be a statistically significant difference. The p-value is the proportion of bootstrap different estimates that are larger than the empirical difference between groups (or smaller, if the difference is negative). Whenever comparing more than two groups, Bonferroni’s correction for multiple comparisons was applied.

- Mixed-effects models with Geisser-Greenhouse correction for matched values were used to compare mean values by mouse across imaging days. Sidak’s multiple comparisons test was used to compare groups on each imaging day.
- Nonparametric one sample Wilcoxon tests were used to determine if group medians were significantly above chance levels in behavior experiments.
- A two-sided Chi squared test with 95% confidence interval was used to compare pass/fail rates in behavior experiments.

## Figure legends

**Figure S1. Velocity and laps run during two photon imaging**. A) Average velocity of mice walking along treadmill belt. B) Average number of laps run per imaging session. Statistics done with mixed effects model. Each point represent an individual animal. Bars represent mean +/- SEM. ns=not significant.

**Figure S2. No difference in active cell number between young and aged mice**. A) Percentage of cells active out of the total cells in a field of view, by mouse. Statistics done with mixed effects model. Bars represent mean +/- SEM. ns=not significant. B) Correlation of percentage active cells versus the total number of cells pooled from all imaging days. Data are shown by mouse.

**Figure S3. Aged mice show a deficit in a spatial memory task**. A) Schematic of the object placement behavioral paradigm. B) Novelty preference of mice measured by percentage of total time spent exploring the novel object. C) Discrimination index calculated by dividing the difference in exploration time between objects by the total exploration time [(novel-familiar)/(novel+familiar)].Pass/fail rate of task, D) where passing was defined as at least a 53% preference for novel object. E) Exploration time dedicated to each object. Statistics in B and C were done with one sample Wilcoxon tests, D with Chi squared test and E with mixed effects model. Bars represent mean +/- SEM. Young N=13, Aged N=13. ns=not significant, *p<0.05, **p<0.01.

