## Supplemental Figures for "Delayed formation of neural representations of space in aged mice"

**Figure S1. Velocity and laps run during two photon imaging.** A) Average velocity of mice walking along treadmill belt. B) Average number of laps run per imaging session. Statistics done with mixed effects model. Each point represent an individual animal. Bars represent mean +/- SEM. ns=not significant.

**Figure S2. No difference in active cell number between young and aged mice.** A) Percentage of cells active out of the total cells in a field of view, by mouse. Statistics done with mixed effects model. Bars represent mean +/- SEM. ns=not significant. B) Correlation of percentage active cells versus the total number of cells pooled from all imaging days. Data are shown by mouse.

**Figure S3. Aged mice show a deficit in a spatial memory task.** A) Schematic of the object placement behavioral paradigm. B) Novelty preference of mice measured by percentage of total time spent exploring the novel object. C) Discrimination index calculated by dividing the difference in exploration time between objects by the total exploration time [(novel-familiar)/(novel+familiar)]. D) Pass/fail rate of task, where passing was defined as at least a 53% preference for novel object. E) Exploration time dedicated to each object. Statistics in B and C were done with one sample Wilcoxon tests, D with Chi squared test and E with mixed effects model. Bars represent mean +/- SEM. Young N=13, Aged N=13. ns=not significant, *p<0.05, **p<0.01.

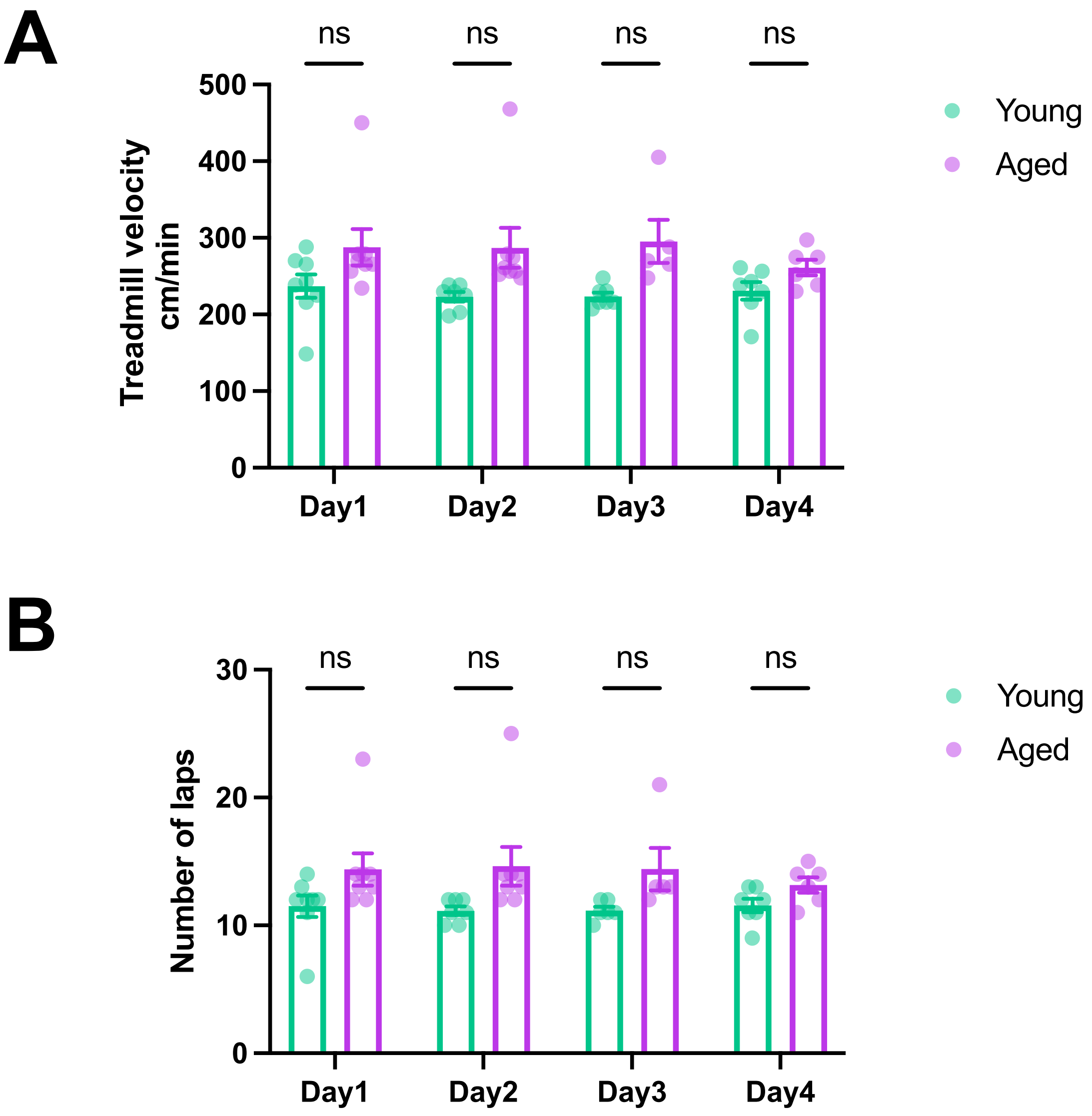
A
ns
ns
ns
ns
500
400
300
200
100
0
Treadmill velocity cm/min
Day1
Day2
Day3
Day4
Young
Aged
B
ns
ns
ns
ns
30
20
10
0
Number of laps
Day1
Day2
Day3
Day4
Young
Aged

### Figure S1

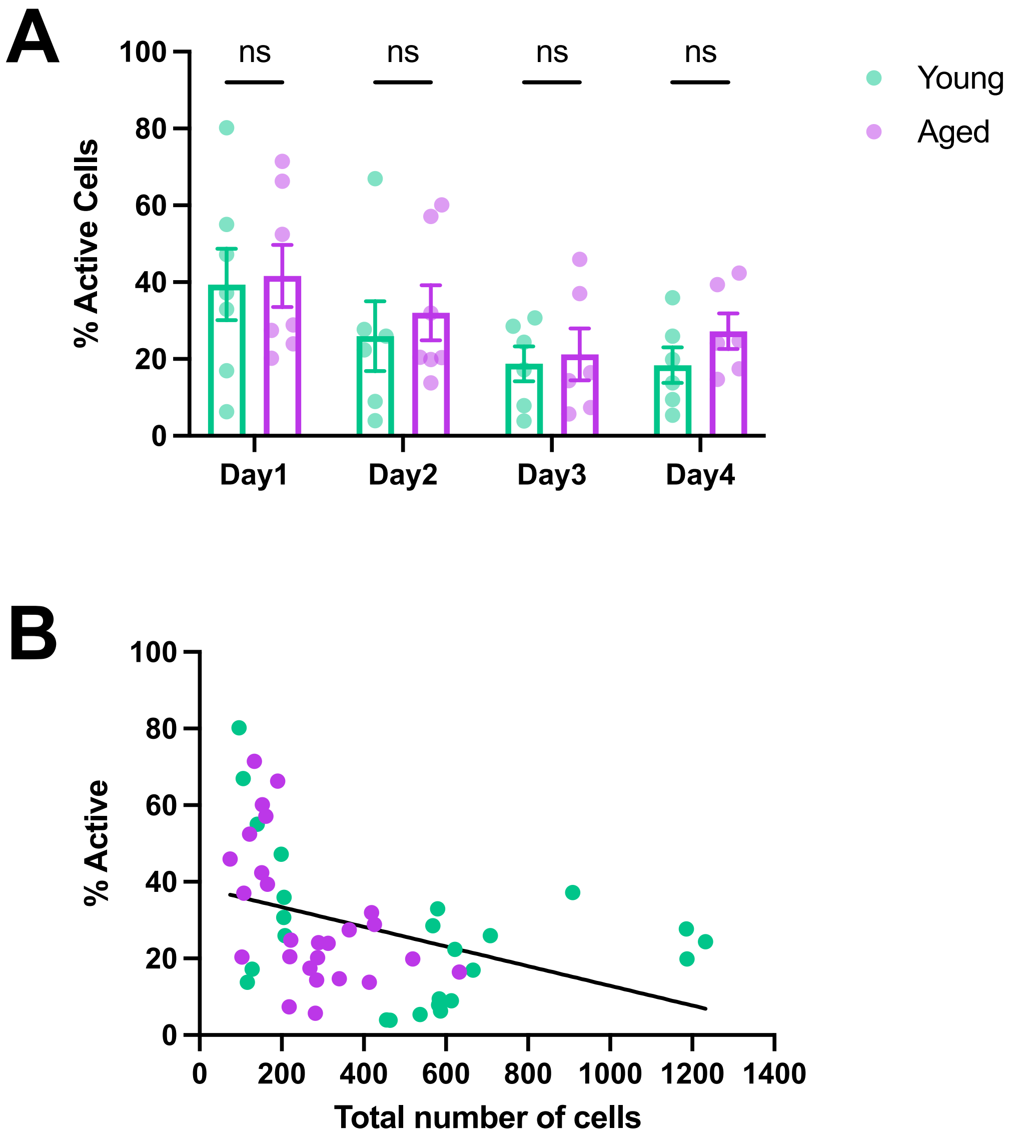
A
100
80
60
40
20
0
% Active Cells
ns
ns
ns
ns
Day1
Day2
Day3
Day4
Young
Aged
B
100
80
60
40
20
0
% Active
0
200
400
600
800
1000
1200
1400
Total number of cells

### Figure S2

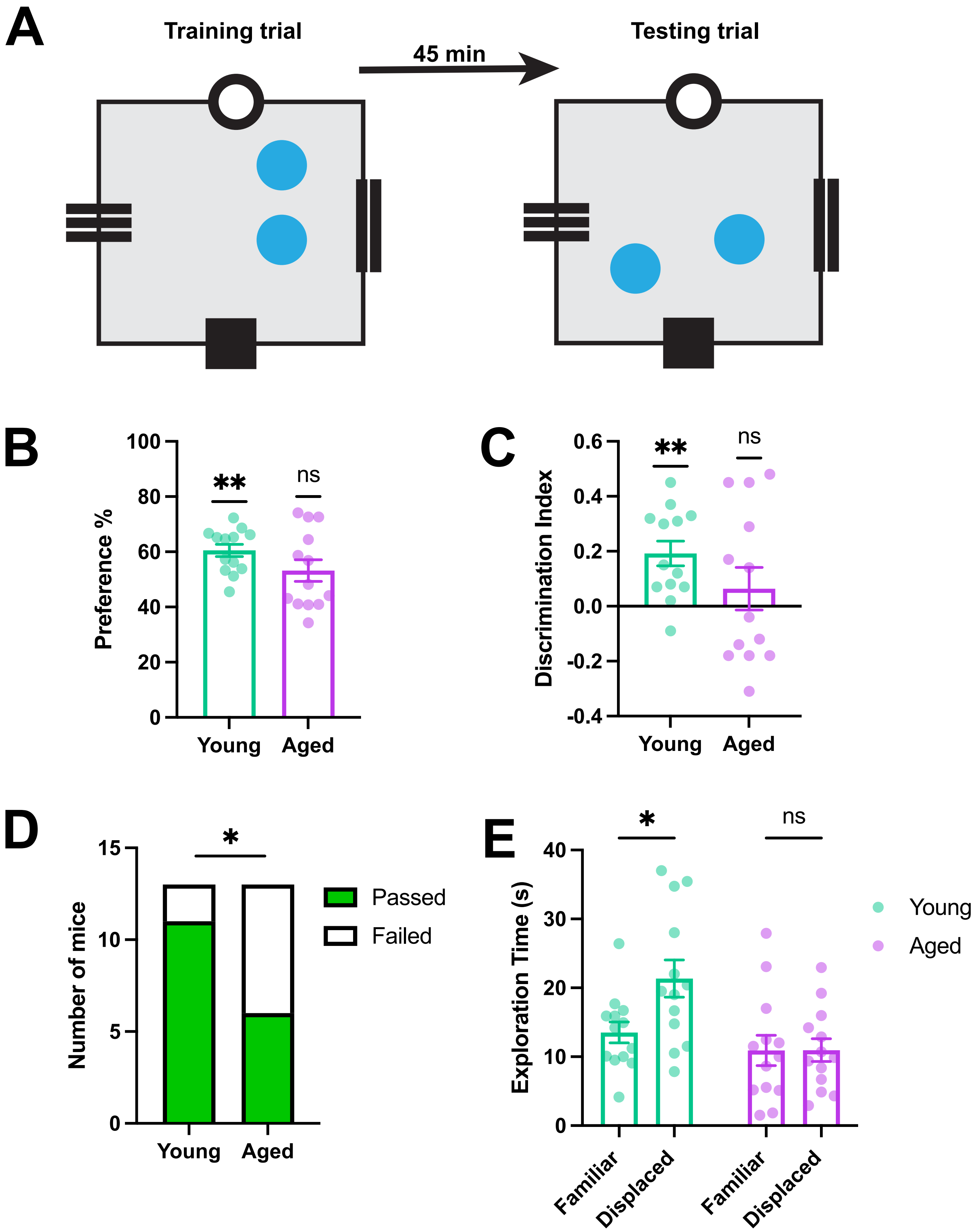

A
Training trial
45 min
Testing trial
B
100
80
60
40
20
0
Preference %
**
ns
Young
Aged
C
0.6
0.4
0.2
0.0
-0.2
-0.4
Discrimination Index
**
ns
Young
Aged
D
15
10
5
0
Number of mice
*
Passed
Failed
Young
Aged
E
40
30
20
10
0
Exploration Time (s)
*
ns
Young
Aged
Familiar
Displaced
Familiar
Displaced


### Figure S3
